# Early detection of sublexical and lexical processing in beginning readers: evidence from Steady-State Visual Evoked Potentials (SSVEPs)

**DOI:** 10.1101/2021.12.07.471641

**Authors:** Fang Wang, Quynh Trang H. Nguyen, Blair Kaneshiro, Lindsey Hasak, Angie M. Wang, Elizabeth Y. Toomarian, Anthony M. Norcia, Bruce D. McCandliss

## Abstract

There are multiple levels of processing relevant to reading that vary in their visual, sublexical and lexical orthographic processing demands. Segregating distinct cortical sources for each of these levels has been challenging in EEG studies of early readers. To address this challenge, we applied recent advances in analyzing high-density EEG using Steady-State Visual Evoked Potentials (SSVEPs) via data-driven Reliable Components Analysis (RCA) in a group of early readers spanning from kindergarten to second grade. Three controlled stimulus contrasts—familiar words versus unfamiliar pseudofonts, familiar words versus orthographically legal pseudowords, and orthographically legal pseudowords versus orthographically illegal nonwords—were used to isolate visual print/letter selectivity, sublexical processing, and lexical processing, respectively. We found robust responses specific to each of these processing levels, even in kindergarteners who have limited knowledge of print. Moreover, comparing amplitudes of these three stimulus contrasts across three reading fluency-based groups and three grade-based groups revealed fluency group and grade group main effects only for lexical contrast (i.e., words versus orthographically legal pseudowords). Furthermore, we found that sublexical orthography-related responses shifted their topographic distribution from the right to left hemisphere from kindergarten to first and second grades. Results suggest that, with more sensitive measures, the sublexical and lexical fine tuning for words—as a bio-marker of reading ability—can be detected at a much earlier stage than previously assumed.

**Declarations of interest:** None

## 1 Introduction

Reading acquisition is a major landmark in child development. In order for children to become fluent readers, they need to build a special form of visual expertise (distinct from expertise in processing objects and faces) that allows them to process words efficiently (McCandliss et al., 2003). Starting with the visual features of single letters, children need to learn correspondences between letters and speech sounds, then learn recurring letter combinations/sub-strings, and finally small words (Dehaene et al., 2005)—a process of retrieving, at minimum, visual word/letter form, sublexical, and lexical information. The current study systematically examines how neural correlates of visual sensitivity for print, sublexical orthographic, and lexical processing manifest and develop in beginning readers.

Starting at the lowest level of the processing hierarchy, visual sensitivity for print—as operationalized by contrasts between words/letter strings and letter-like objects (e.g., symbols, pseudofonts)—is also referred to as “coarse neural tuning for print” or “letter selectivity” (Maurer, Brandeis, & McCandliss, 2005; Maurer et al., 2006; Wong et al., 2005). Coarse neural tuning for print has been extensively investigated in different writing systems (alphabetic: Brem et al. (2005); Maurer, Brandeis, & McCandliss (2005); Zhao et al. (2014); Eberhard-Moscicka et al. (2015, 2016); logographic: Maurer et al. (2008); Okumura et al. (2014); Tong et al. (2016); Wang & Maurer (2017); Zhao et al. (2012, 2019)), in adults (Bentin et al., 1999; Brem et al., 2005), in children of different age groups (Maurer et al., 2006; Posner & McCandliss, 1999), as well as in atypical readers (Brem et al., 2020; Mahé et al., 2012; Maurer & McCandliss, 2007; Maurer et al., 2011; Pleisch et al., 2019).

Coarse print tuning is typically reflected by increased neural activity to words compared to letter-like objects (e.g., pseudofonts) (Maurer, Brem, et al., 2005; Centanni et al., 2017, 2018, 2019). Developmentally, coarse tuning starts to emerge when children are beginning to read, following an inverted U-curve, with an initial increase and then a later decrease starting in second grade (Maurer et al., 2006, 2011).

Numerous functional magnetic resonance imaging (fMRI) studies have localized an area in the left ventral occipito-temporal (vOT) cortex that exhibits coarse print tuning (McCandliss et al., 2003; Cohen et al., 2000; Dehaene & Cohen, 2011). High temporal resolution electroencephalogram (EEG) studies have revealed that coarse tuning occurs 120 to 280 ms after stimulus onset over left occipito-temporal areas (Maurer et al., 2006). However, more recent studies have found that visual word recognition is not only limited to the left vOT, but also involves other language areas (e.g., midfusiform sulcus, angular gyrus) that are responsible for sublexical and lexical information processing (with fMRI: Bouhali et al. (2019); Lerma-Usabiaga et al. (2018); White et al. (2019); with intracranial recording: Woolnough et al. (2021)). This claim has been further supported by our recent EEG study with adult participants (Wang et al., 2021) that used a data-driven spatial filtering approach—Reliable Components Analysis (RCA) (Dmochowski et al., 2012, 2015)—which could successfully distinguish different processes having overlapping time courses and interacting topographic patterns. In our previous study, distinct neural sources—one being maximal over left vOT regions, and the other being maximal over more dorsal parietal regions—were found when contrasting visual words with pseudofont visual controls (Wang et al., 2021). Using the same approach, the present study asked whether multiple sources are also involved in coarse print tuning in beginning readers, starting as early as children in kindergarten.

Contrasts between words and pseudofonts/symbols mix information from visual letter forms, sublexical orthographic structures, and lexical properties. This raises the questions of which specific properties of words trigger coarse print tuning measured with words-pseudofonts contrast? To what extent is the visual specialization for words sensitive to sublexical and/or lexical properties? More importantly, how do these functionally distinct yet likely temporally overlapping levels of processing develop during early reading acquisition? Therefore, in addition to the development of coarse tuning, the current study also systematically isolated sublexical and lexical processing using specifically-designed stimulus contrasts to address such questions.

In previous studies, sublexical and lexical level orthographic processing have been referred to as “fine neural tuning for word” (Maurer, Brandeis, & McCandliss, 2005; Maurer et al., 2006; Wong et al., 2005; Zhao et al., 2014). Specifically, sublexical tuning reflects sensitivity to sequential dependencies and letter position frequencies (orthographically legal pseudowords vs. orthographically illegal nonwords)—orthographic familiarity; lexical tuning is associated with sensitivity to well-formed letter arrays that define printed words (words vs. pseudowords)—lexicality (Araújo et al., 2015).

With regards to lexical fine tuning, inconsistent results have been found across studies and scripts. One set of results has demonstrated stronger neural responses to words than to word-like stimuli. Other studies, however, revealed the opposite direction (i.e., word-like stimuli > words, Brem et al. (2009)) or even null effects (Araújo et al., 2012; Eberhard-Moscicka et al., 2015; Hauk et al., 2006; Maurer, Brandeis, & McCandliss, 2005; Proverbio et al., 2004; Zhao et al., 2014). For instance, larger ERP amplitudes for words over pseudowords have been reported in German-speaking second graders (Maurer et al., 2006) but not in first graders (Eberhard-Moscicka et al., 2015; Zhao et al., 2014), adults, or kindergarteners (Maurer, Brem, et al., 2005). By contrast, some studies with adults have found larger activity for pseudowords than for words (for example, with EEG: Hauk et al. (2006)). Moreover, findings on when fine tuning emerges during development are also mixed. For example, Posner & McCandliss reported fine tuning in 10-year-olds, but not in 7- and 4-year-old English speaking children (Posner & McCandliss, 1999). Some German studies detected fine tuning in a group of 8.26-year-old children (Maurer et al., 2006), while another German study found fine tuning only in 7-year-old children with high reading ability but not in those with low reading ability (Zhao et al., 2014). It has been suggested that large individual differences in young children’s reading ability (Jenkins et al., 2003) are an important factor that affects tracking the emergence of fine tuning (Eberhard-Moscicka et al., 2015; Zhao et al., 2014), especially because the effect size of fine tuning is rather small when compared with the effect size of coarse tuning.

Due to the difficulty of isolating discriminative responses between largely overlapped visual stimuli (e.g., words vs. pseudowords) in beginning readers, more sensitive and specific assays are required. Recently, Lochy and colleagues have investigated lexical fine tuning using the contrast of words versus pseudowords with a sensitive steady-state visual evoked potentials (SSVEP) paradigm. Following the “base/deviant” stimulation mode, the SSVEP paradigm presents a sequence of stimuli at two distinct and predefined periodic rates, for example, 2 Hz (500 ms per item) word deviants were embedded in a stream of pseudowords to form a 10 Hz (100 ms per item) base stimulation frequency. Periodic responses to base and deviants are elicited at the predefined stimulation frequency and its harmonics (i.e., integer multiples of the stimulation frequency). Due to it’s small noise bandwidth, this paradigm can provide high signal-to-noise (SNR) ratio in only a few minutes (Lochy et al., 2015, 2016, 2020). In one of their EEG studies with adults, larger responses to words than to pseudowords were robustly detected in every participant. However, when testing the same contrast (i.e., words vs. pseudowords) in preschoolers and children at first and second grade, no evidence of discrimination between these two types of stimuli was found (Lochy et al., 2016; van de Walle de Ghelcke et al., 2021).

The presentation rates used by Lochy et al. (2016) for children were 1.2 Hz and 6 Hz for deviant and base stimulation, respectively. These stimulation rates might be too high for beginning readers, especially when it comes to demanding, lexical-level processing. This conjecture is based on our recent finding that in adults, faster presentation rates evoked weaker responses compared to slower rates (Wang et al., 2021). In addition, we showed that “image alternation” mode, comprising a periodic base rate with every other image being a deviant (e.g., 6/2=3 Hz) elicits responses with higher SNR, compared with the mode wherein deviant stimuli are presented at a sub-multiple greater than twice (e.g., 10/5=2 Hz) the base rate. Furthermore, Yeatman & Norcia (2016) found that the best temporal frequency for driving cortical responses for text is 1 Hz, substantially lower than that for faces (4 Hz). Therefore, the present study used the words-pseudowords contrast and SSVEP paradigm, but slowed down the stimulation rates to 1/2 Hz in an alternation mode. In addition, instead of only focusing on several pre-selected sensors in the left hemisphere as is done in most previous SSVEP work (Lochy et al., 2015, 2016, 2020), we used a spatial filtering approach (RCA) that can recover multiple underlying neural sources that have partially overlapping topographies (Wang et al., 2021). Finally, rather than using a passive viewing task, we used a repetition detection task to keep children more engaged and thus potentially more able to engage in higher levels of visual processing demands, which has been shown to be important for observing activity related to higher-level of reading-related processes (e.g., lexicality effects) (Yang et al., 2012).

As compared to the literature on coarse and lexical fine tuning, research on sublexical orthography processing is less well-developed, even though orthography perception is a prerequisite for word perception (Cheng & Caldwell-Harris, 2010). To our knowledge, no studies have examined the development of sublexical tuning in very beginning readers. Previous studies on adults have been limited to examining orthographic regularity by comparing words with typical vs. atypical orthography (different bigram or trigram frequency of letter combinations) (Binder et al., 2006; Woollams et al., 2011). However, manipulation of orthographic regularity within words may recruit additional processes beyond orthographic processing (e.g., familiarity or meaningfulness) due to the tight coupling between orthography and the lexicon in real words (Zhao et al., 2019). Therefore, it has been suggested that the contrast between pseudowords and consonant strings can better isolate sublexical orthographic regularity (Posner & McCandliss, 1999). One EEG study with 7-year-olds found larger responses to pseudowords relative to consonant strings only in children with higher reading ability (Zhao et al., 2014). It is thus possible that different levels of reading expertise may affect the emergence and developmental trajectory of fine tuning for sublexical orthographic patterns. Instead of pure consonant strings, the current study used orthographically illegal nonwords created by reordering letters that appeared in the pseudowords (CVC structure), an approach that avoids potential confounding of consonant-vowel differences. More importantly, we recruited kindergarteners, first and second graders to investigate the developmental changes of this effect.

To summarize, the current study systematically manipulated stimulus contrasts to isolate letter/word form, sublexical, and lexical variants of processing during early reading acquisition. In order to track the early development changes of these different levels of processing, SSVEP-EEG data were recorded with slower stimulation rates (i.e., 1/2 Hz) from kindergarteners up to second graders in a relatively demanding repetition detection task. To capture multiple underlying temporal/topographic sources, a spatial filtering technique—RCA—was employed.

## 2 Methods

### 2.1 Ethics Statement

The study was approved by the Institutional Review Board of Stanford University. A parent or legal guardian of each participant received a written description of the study and gave written informed consent before the beginning of the session; each participant also assented to participating.

### 2.2 Participants

A total of 57 healthy, English-speaking children with normal or corrected-to-normal vision and who had no reading disabilities participated. Out of these initial 57 subjects, nine were excluded from further analyses due to having withdrawn after behavioral assessments (*N* = 2) or due to EEG data quality issues (i.e., excessive movement during recording, incomplete EEG data, *N* = 4) or low performance in the repetition detection task during EEG recording (*N* = 3). The remaining sample (*N* = 48), whose EEG data were analyzed, included 15 kindergarteners (11 males, *m* = 5.9 years, *s* = 0.4 years), 16 first graders (9 males, *m* = 6.9 years, *s* = 0.3 years, and 17 second graders (7 males, *m* = 7.6 years, *s* = 0.3 years).

### 2.3 Behavioral Assessment

Behavioral assessments were administered on average 8.5 days (*s* = 5.2 days) before the EEG session. We tested children on two sub-tests of the Comprehensive Test of Phonological Processing, Second Edition (CTOPP-II, Wagner et al. (2013)) to assess their phonological awareness and rapid naming abilities. We tested children’s word recognition and decoding using the Test of Word Reading Efficiency, Second Edition, (TOWRE-II, Torgesen et al. (2012)), and letter-word identification subset of Woodcock-Johnson Tests of Achievement, Fourth Edition (WJ-IV; Schrank et al. (2014)), respectively.

### 2.4 Stimuli

The study involved four types of stimuli: English words (W), pseudofonts (PF), orthographically legal psuedowords (OLPW), and orthographically illegal nonwords (OINW). All stimuli were composed of 3 letters or pseudoletters. W stimuli were high-frequency words (mean 2110 per million, range 629–5851 per million) with consonant-vowel-consonant (CVC) structure. W, OLPW, and OINW stimuli were rendered in Courier New font, while PF stimuli were rendered from the Brussels Artificial Character Set font (BACS-2, Vidal & Chetail (2017)); here, Courier New glyphs composing the English words were mapped to corresponding pseudofont glyphs. OLPW were built on an item-by-item basis by exchanging letters across intact words while retaining the CVC structure. OLPW were thus pronounceable and well-matched for orthographic properties of intact words. OINW were also built on an item-by-item basis by shuffling the letters across the set of OLPW, resulting in unpronounceable, statistically implausible letter-string combinations. Bigram frequencies were well matched between words and orthographically legal pseudowords (*t*(30) = 0.26, *p* = 0.79), while bigram frequencies of orthographically legal pseudowords and orthographically illegal nonwords were significantly different (*t*(30) = 6.26, *p* < 0.001). In all, the stimuli set comprised 32 high-frequency words (W), 16 pseudofont strings (PF), 32 orthographically legal pseudowords (OLPW), and 16 orthographically illegal nonwords (OINW), for a total of 96 stimulus exemplars.

Three experimental conditions were investigated. Condition 1 involved words alternating with pseudofonts (W–PF). This condition was designed to probe coarse print tuning. Condition 2 alternated words with orthographically legal pseudowords (W–OLPW) and was designed to probe lexical fine tuning (lexicality effect). In condition 3, orthographically legal pseudowords were alternated with orthographically illegal nonwords (OLPW–OINW) in order to probe sublexical fine tuning (orthographic regularity effect). In each condition, the *base* rate of visual stimulation was 2 Hz, while stimuli from a given category appeared in alternation at half that rate (i.e., at a *deviant* rate of 1 Hz). The stimulus conditions are summarized in Table 1, and examples are shown in Figure 1.

**Table 1:** Stimulus conditions. Conditions 1 and 2 assessed processing of word deviants relative to pseudofonts (W–PF) and orthographically legal pseudowords (W–OLPW), respectively. Condition 3 assessed processing of orthographically legal pseudowords relative to orthographically illegal nonwords (OLPW–OINW). All contrasts were presented centered on the screen with frequency rates of 1 Hz deviant and 2 Hz base.

|  | Condition 1 | Condition 2 | Condition 3 |
| --- | --- | --- | --- |
| **Stimuli** | W–PF | W–OLPW | OLPW–OINW |
| **Deviant / Base** | 1 Hz / 2 Hz | 1 Hz / 2 Hz | 1 Hz / 2 Hz |

**Figure 1:**
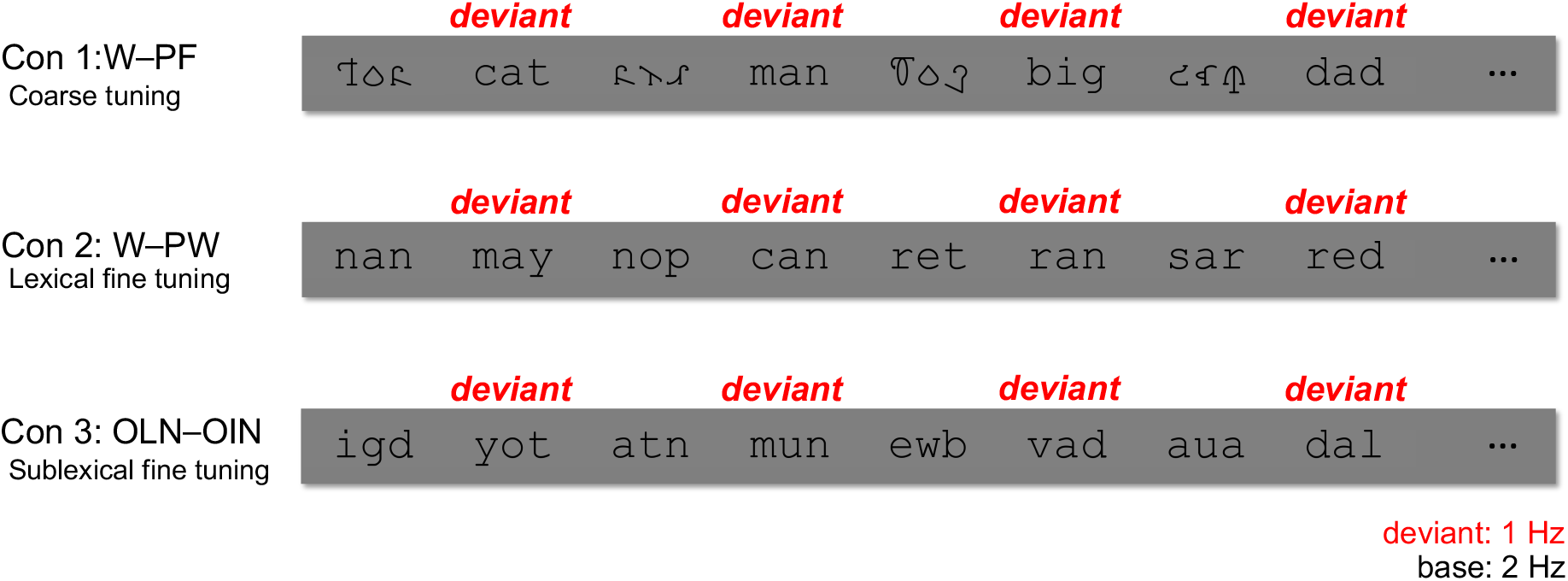
Experimental Design. Examples of stimuli presented in the experiment. 1 Hz deviants were embedded within a 2 Hz base stream in all three conditions. The first condition assessed coarse print tuning with words alternating with pseudofonts (W–PF). The second condition assessed lexical fine tuning with words alternating with /orthographically legal pseudowords (W–OLPW). The third condition assessed sublexical fine tuning with orthographically legal pseudowords alternating with orthographically illegal nonwords (OLPW–OINW).

### 2.5 Experimental Procedure

The three conditions were presented in a fixed order (condition 1 followed by condition 2 and then condition 3). Using in-house stimulus presentation software, each condition started with a blank screen, of which the duration was jittered in the range of 1500 ms to 2500 ms. Then, a given image was presented for 500 ms and a complete cycle of alternation lasted 1000 ms; these periods result in what we refer to as a 2 Hz base frequency and 1 Hz deviant frequency, respectively. All stimuli were presented at the center of the screen in black font on a gray background. The stimulus images were 600 × 160 pixels in size, spanning 7.5 (horizontal) by 2 (vertical) degrees of visual angle. Participants were asked to press a button on an external response pad with their preferred hand when a stimulus (i.e., target) was repeated three times in a row. Among the 12 trials (each trial lasted for 12 seconds) that were pseudorandomly presented for each condition, there were four “non-target” trials, which contained no repeated stimuli (Figure 2A); four “terminal” trials in which repeated stimuli appeared at the end (Figure 2B); and four “catch” trials with repeated stimuli randomly appearing elsewhere during the trial (Figure 2C). Each terminal and catch trial contained only one target. Participants were given verbal feedback about their performance after the end of each trial. Due to excessive movements from the button press, EEG data of the entire durations (i.e., 12 seconds) of the four catch trials were excluded from further analysis for each participant. Data corresponding to the four terminal trials were still included because the movements happened at the end of the trial. Participants were allowed to take short breaks between trials and conditions as needed.

**Figure 2:**
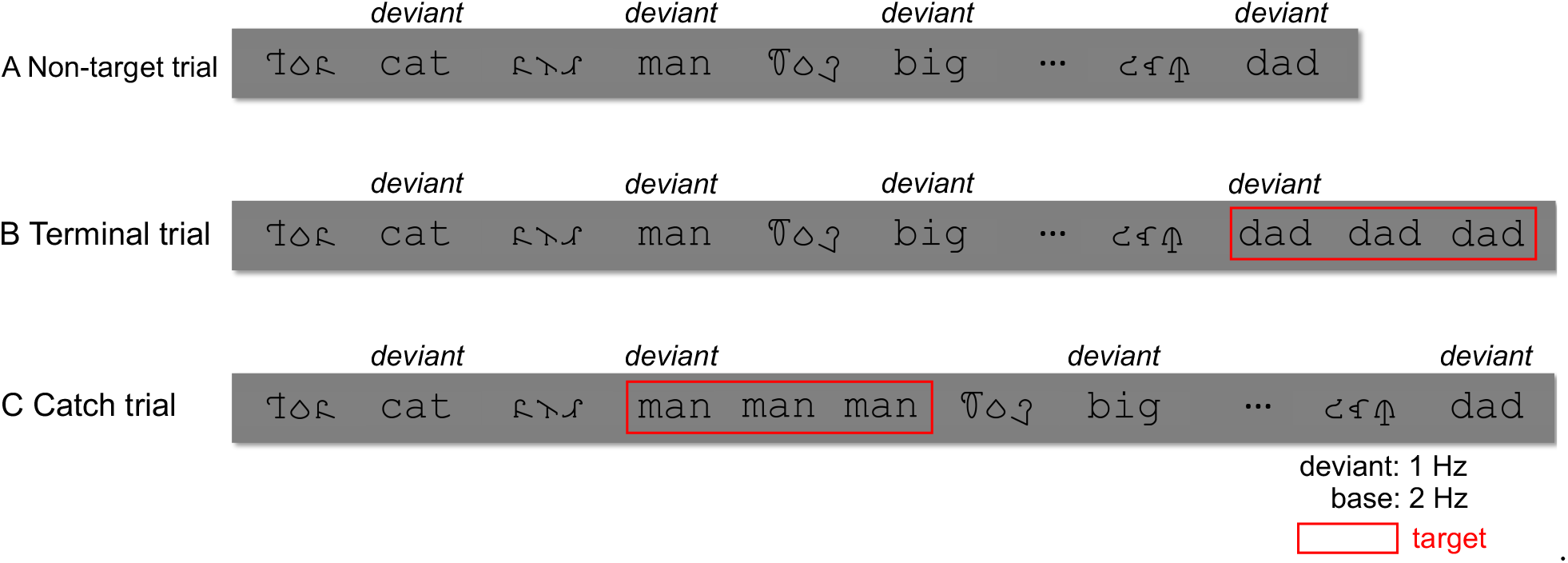
Illustration of different types of trials in condition 1: W–PF. Twelve trials were pseudorandomly presented for each condition (herein only take condition 1 for example), including four non-target trials, four terminal trials, and four catch trials. Every participant’s data corresponding to the four catch trials were excluded from EEG analyses due to excessive response-related movements during recording.

Participants were seated 1 m away from the computer screen in a dimly lit room. Prior to the EEG recording, the experimenter led the participants through a brief practice session to familiarize them with the 3-in-a-row repetition detection task and experimental procedure.

As compensation, each child received a sticker after the study. The entire experiment lasted about 40 minutes including setup, practice, and breaks.

### 2.6 EEG Recording and Preprocessing

128-channel EEG were recorded with the Electrical Geodesics, Inc. (EGI) system (Tucker, 1993), using a Net Amps 300 amplifier and geodesic sensor net against Cz (vertex) reference. Data were sampled at 500 Hz. Impedances were kept below 50 kΩ. After recording, the data were bandpass filtered offline (zero-phase filter, 0.3–50 Hz) using Net Station Waveform Tools.

Filtered data were then imported into an in-house signal processing software for preprocessing. EEG data were first re-sampled to 420 Hz to ensure an integer number (herein 7) of time samples per video frame at a frame rate of 60 Hz, as well as integer numbers of frames (i.e., 60 frames per deviant cycle and 30 frames per base cycle) per cycle for the current stimulation frequencies (1 Hz deviant and 2 Hz base). Then, sensors for which more than 15 % of samples exceeded a ± 30 *μV* amplitude threshold were interpolated by averaging data from six surrounding sensors. The continuous EEG was filtered using Recursive Least Squares (RLS) filters (Tang & Norcia, 1995) and re-referenced to average reference Lehmann & Skrandies (1980). Re-referenced data were segmented into 1-second epochs. Epochs with more than 10 % of time samples exceeding a ± 30 *μV* noise threshold, or with any time sample exceeding an artifact threshold of (± 60 *μV*) (reflecting e.g., eye blinks, eye movements or body movements), were rejected from further analyses. The RLS filters were tuned to each of the analysis frequencies (i.e., base frequency, deviant frequency, and their harmonics) and converted to complex values by means of the Fourier transform. Complex-valued RLS outputs were decomposed into real and imaginary Fourier coefficients as input for the spatial filtering computations as described below.

### 2.7 Analysis of EEG Data

#### 2.7.1 Reliable Components Analysis (RCA)

Reliable Components Analysis (RCA) is a matrix decomposition technique that derives a set of spatial reliable components (RCs) from the 128-sensor array by computing optimal weightings of electrodes to maximize between-trial covariance relative to within-trial covariance (Dmochowski et al., 2012, 2015; Wang et al., 2021). Since response phases of SSVEP are constant over repeated stimulations, the activations of each component reflect phase-locked activities. RCA is an eigenvalue decomposition technique; the eigenvectors represent linear weightings of sensors (i.e., linear spatial filters) by which the electrode-space data matrix is projected to a component-space matrix. The resulting spatial component data exhibit maximal Pearson Product Moment Correlation Coefficients (Pearson, 1896) across trials. The percentage of “reliability” explained by individual components are computed from the corresponding eigenvalues (Dmochowski et al., 2015). This eigenvalue decomposition procedure returns RCs sorted in descending order of reliability explained; that is, the first component (RC1) explains the most reliability in the data. Each RC can be visualized as scalp topographies using a forward-model projection of the spatial filter weight vectors (Parra et al., 2005). More details on the spatial filtering technique are provided by Dmochowski et al. (2012, 2015).

#### 2.7.2 RCA at deviant frequency and its harmonics

In order to investigate the discriminative responses between deviant and control stimuli (W– PF, W–OLPW, and OLPW–OINW), we first computed RCA over real and imaginary Fourier coefficients at the deviant frequency and its first five odd harmonics (excluding even—herein base—harmonics): 1 Hz, 3 Hz, 5 Hz, 7 Hz, and 9 Hz (excluding 2 Hz, 4 Hz, 6 Hz, 8 Hz, and 10 Hz). In order to assess processing of different levels of information during visual word recognition (i.e., letter/word selectivity, lexicality, and sublexical orthographic regularity, respectively, in conditions 1, 2, and 3), we computed RCA separately for each condition.

#### 2.7.3 RCA at base frequency and its harmonics

In order to test whether visual responses to low-level stimulus features were comparable across conditions, as well as to explore potential underlying sources in a data-driven fashion instead of picking several electrodes of interest a priori (e.g., three middle occipital electrodes) as in most of previous SSVEP work (Lochy et al., 2015, 2016, 2020; van de Walle de Ghelcke et al., 2021), we also computed RCA at the base frequency and its first five harmonics (i.e, 2 Hz, 4 Hz, 6 Hz, 8 Hz, and 10 Hz) over the 128-sensor array. To enable a direct quantitative comparison of the three conditions in a shared component space, we computed the RCA weights over the three conditions together.

#### 2.7.4 Analysis of Component-Space Data

For RCA at the deviant frequency and its harmonics (1 Hz, 3 Hz, 5 Hz, 7 Hz, and 9 Hz), statistical analyses were performed on the projected data of the first three reliable components returned by RCA. We report results from all three RCs for condition 1 (W–PF), and from the first RC for conditions 2 (W–OLPW) and 3 (OLPW–OINW), as these were the components associated with statistically significant component-space data.

For RCA at the base frequency and its harmonics (2 Hz, 4 Hz, 6 Hz, 8 Hz, and 10 Hz), given the substantial percentage of reliability each RC explained in the data (as computed from the RCA eigenvalues (Dmochowski et al., 2015); details reported in Results), statistical analyses were performed on the first three reliable components.

For both the deviant and base analyses, we first projected the sensor-space data through each spatial filter weight vector for each component. Then, the projected data were averaged across 1-second epochs on a per-participant basis. Finally, statistical analyses were performed across the distribution of all participants. Statistical significance of component-space responses was determined by a Hotelling’s two-sample t^2^ test (Victor & Mast, 1991) on the distribution of real and imaginary Fourier coefficients at each harmonic in each component. Multiple comparisons were corrected using False Discovery Rate (FDR; Benjamini & Yekutieli (2001)). Fifteen comparisons (5 harmonics × 3 components) were corrected for base analyses involving three conditions together and for separate deviant analyses in conditions 1, 2 and 3, respectively.

For deviant RCA analyses in coarse print tuning (condition 1: W–PF, see Figure 3), we first present topographies of the spatially filtered components (Figure 3A). Next, mean responses of projected data are visualized as vectors in the 2D complex plane, with the length of the vector representing the response amplitude and the angle of the vector relative to 0 degree (counterclockwise from the 3 o’clock direction) representing the phase. The error ellipses around the tips of vectors are the standard errors of the mean (*SEM*). Finally, amplitudes (*μV*) at each harmonic in each component were plotted in bar plots, with significant responses (according to adjusted *p_FDR_* values of t^2^ tests of the complex data) indicated with asterisks (Figure 3C). Phase values (degrees) were plotted across harmonics. We additionally plotted a line of best fit through the significant harmonics, as each component contained statistically significant activity at at least two harmonics (Figure 3C). Latencies (in ms) were estimated by the slope of this line of best fit (Norcia et al., 2020).

**Figure 3:**
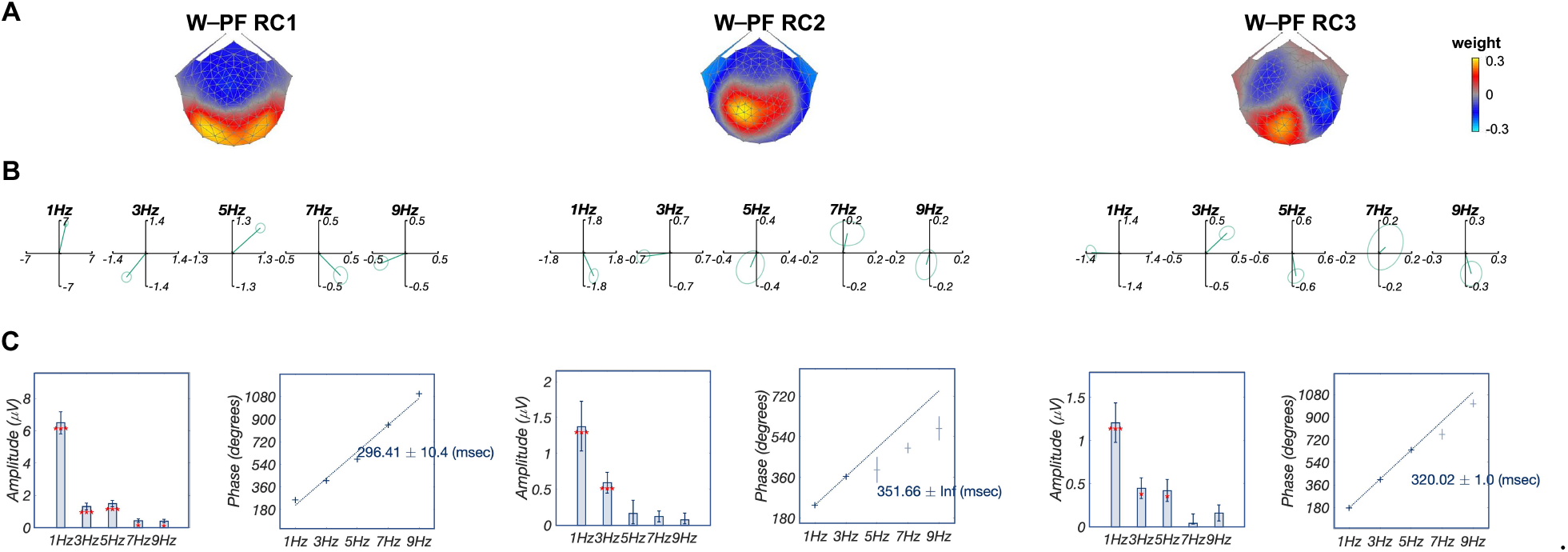
Deviant Analysis: multiple sources underlying coarse print tuning (condition 1: W–PF) in beginning readers. A: Topographic visualizations of the spatial filters for the first three components: RC1, RC2, and RC3; B: Response data in the complex plane, where amplitude information is represented as the length of the vectors, and phase information in the angle of the vector relative to 0 degrees (counterclockwise from 3 o’clock direction), ellipse indicates standard error of the mean for both amplitude and phase; C: Projected amplitude (left) for each harmonic in bar charts, *: *p_FDR_* < 0.05, ***: *p_FDR_* < 0.001, as well as the latency estimation computed from the slope of the line fit across significant harmonics for each component: 296.41 ms, 351.66 ms, and 320.02 ms respectively for components 1–3.

For deviant RCA analyses in lexical (condition 2: W–OLPW) and sublexical (condition 3: OLPW–OINW) fine tuning (see Figure 4), besides topographies (Figure 4A&B) and amplitude bar plots (Figure 4D), we present mean responses of projected data at 1 Hz—the only significant harmonic in both conditions 2 and 3—as vectors in the 2D complex plane, since the presence of only one significant harmonic in condition 2 prevents us from estimating latency from the phase slope. To examine the time course difference between lexical and sublexical fine tuning, we used the Circular Statistics toolbox (Berens et al., 2009) to compare distributions of RC1 phases at 1 Hz in these two conditions, additionally mean RC1 phase difference is reported in ms.

**Figure 4:**
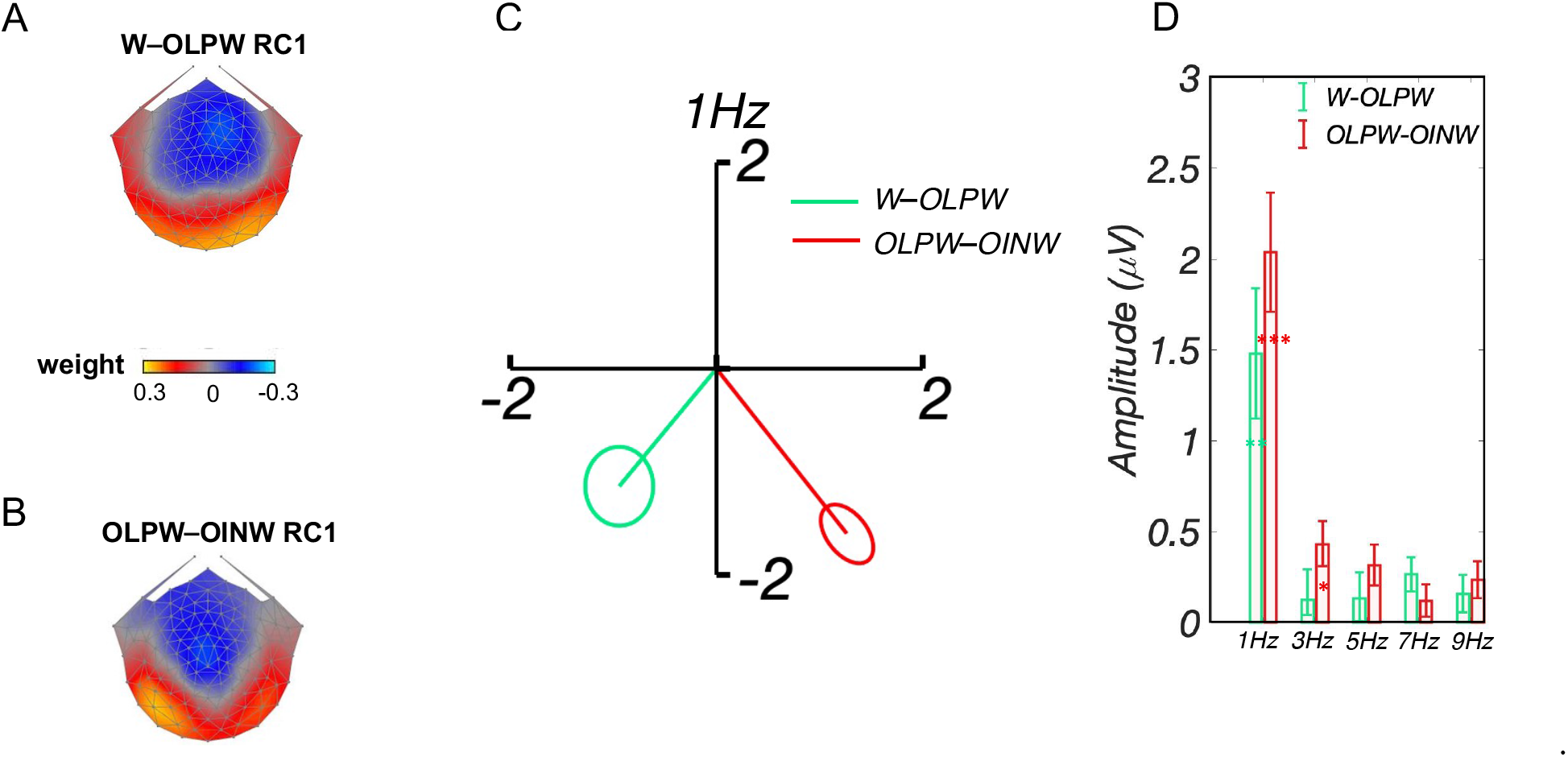
Deviant Analysis: lexical (lexicality effect) and sublexical fine tuning (orthographic regularity effect) in beginning readers. A&B: Topographic visualizations of the spatial filters for the first component (RC1) in W–OLPW (top) and OLPW–OINW (bottom), respectively; C: Response data at the first harmonic (1 Hz, the only significant harmonic in both conditions) presented in the complex plane, where amplitude information is represented as the length of the vector, and phase information in the angle of the vector relative to 0 degree (counterclockwise from 3 o’clock direction), ellipse indicates standard error of the mean (*SEM*). Different phases were found in sublexical and lexical tuning, indicating the two components reflect different processing. D: Bar charts of projected amplitude at each harmonic for RC1, in W–OLPW (green) and OLPW–OINW (red), respectively.

To explore possible brain-behavior relationships, we partitioned the participants into three groups based on grade (grade group: kindergarten (*N* = 15), first grade (*N* = 16), and second grade (*N* = 17)) and three groups based on reading fluency raw scores (measured with TOWRE, reading group: low reading group (*N* = 16), middle reading group (*N* = 16), and high reading group (*N* = 16)). First, to enable a direct quantitative comparison of different groups in a shared component space, projected amplitudes were extracted from deviant RCA output that trained on all subjects together for each condition. One-way ANOVAs were computed separately for each component—the first three components in condition 1, the first component in conditions 2 and 3—using the amplitude at the first harmonic (1 Hz), as this harmonic had the most robust signal in all three conditions. Statistically significant ANOVAs were followed by post hoc pairwise comparisons. In addition, for inspection of differing spatial filter topographies across groups, RCA was also trained separately on each group (both reading and grade) for each condition. Only topographies for conditions with consistently significant component-space activity across three groups—i.e., at least one harmonic of the projected data was significant—are presented in Results.

Finally, for base RCA computed on all three conditions together, as with the deviant RCA described above, we also visualized the projected data in four ways: Spatial filter topographies, mean responses as vectors in the 2D complex plane, amplitude in bar charts, and phase values (degrees) accompanied by a line of best fit and slope per condition.

### 2.8 Behavioral Analysis

For behavioral responses to the 3-in-a-row repetition detection task during EEG recording, we computed *d*’ based on the z-transformed probabilities of hits and false alarms (Macmillan & Creelman, 2004). One-way ANOVA with within-factor of condition was computed on *d*’ across three conditions.

## 3 Results

### 3.1 Deviant Analyses Results

We computed RCA at the deviant stimulation frequency and its harmonics for each condition separately in order to investigate potentially different topographies and response dynamics underlying different levels of information processing in visual word recognition.

For responses to words alternating with pseudofonts (condition 1), the first three reliable components each contain significant driven activity (i.e., at least one harmonic is significant) and together explain 70% of the reliability in the data. As shown in Figure 3A, the topography of the first component (RC1) includes bilateral peaks, but is maximal at left posterior vOT electrodes; the second component (RC2) is displaced to more central-parietal electrodes with left lateralization; and the third component (RC3) has a more complex topography over occipital electrodes. For RC1, component-space data were significant at all five harmonics (1 Hz, 3 Hz, and 5 Hz, *p_FDR_* < 0.001; 7 Hz and 9 Hz, *p_FDR_* < 0.05; corrected for 15 comparisons). Response phase over these harmonics is linear as can be seen in the left panel of Figure 3C, allowing us to estimate response latency from the slope of the function. This procedure produced a latency estimate of 296.41 ± 10.4 ms. For RC2, projected data were significant at the first two harmonics only (1 Hz and 3 Hz, *p_FDR_* < 0.01, corrected for 15 comparisons; Figure 3B, middle). The linear fit through these two harmonics produced a latency estimate of 351.66 ms (Figure 3C, middle; *SEM* is unavailable whenever there are only two significant points). For RC3, projected data were significant over the first three harmonics (1 Hz, *p_FDR_* < 0.001; 3 Hz and 5 Hz, *p_FDR_* < 0.05; corrected for 15 comparisons; Figure 3B, right), with a latency estimate of 320.02 ± 1.0 ms (Figure 3C, right). In all, the responses in each component are dominated by the lowest-frequency odd harmonic, while latency estimates indicate that activity in RC1 leads activity in RC2 and RC3.

For responses to W alternating with OLPW (condition 2) and OLPW alternating with OINW (condition 3), only RC1 contained significant stimulus-driven activity. RC1 in conditions 2 and 3 explained 23% and 31% of the reliability in the data, respectively. RC1 in condition 2 is broadly distributed over occipital-temporal-parietal electrodes (Figure 4A), with only the first harmonic (1 Hz) being statistically significant (*p_FDR_* < 0.01; corrected for 15 comparisons; Figure 4D). RC1 in condition 3 is maximal over occipital electrodes, with a bias to the left hemisphere (Figure 4B). Only the first two harmonics were significant after FDR correction (1 Hz, *p_FDR_* < 0.001; 3 Hz, *p_FDR_* < 0.05; corrected for 15 comparisons; Figure 4D). The comparison of responses at 1 Hz for both conditions in the complex plane (Figure 4C), shows distinct phases (vector angles) between these two conditions. Circular statistics of RC1 phase comparison in these two conditions showed that RC1 phases in condition 3 (OLPW–OINW) are significantly longer (phase difference at 1 Hz: 243.6 ms, *p* < 0.001) than that in condition 2 (W-OLPW). Different phases indicate that conditions 2 and 3 evoked different levels of information processing (i.e., lexical and sublexical processing), with overlapping topographies.

For the exploration of brain-behavior relationships, in terms of response projected amplitude, significant group effects were only found in condition 2 (W–OLPW, lexical fine tuning effect, reading group (*F*(2, 46) = 3.46, *p* < 0.05) and grade group (*F*(2, 46) = 7.28, *p* < 0.01)). To further explore these group effects, post hoc pairwise t tests showed that response amplitudes at the first harmonic (i.e., 1 Hz) were stronger in middle (*p* < 0.05) and high (*p* < 0.05) reading groups than in the low reading group, and no significant difference was found between middle and high groups (*p* = 0.67) (see Figure 5, left). In addition, response amplitudes at the first harmonic were stronger in first grade than in kindergarten (*p_FDR_* < 0.01) and second grade (*p_FDR_* < 0.05), and a marginally significant difference was found between kindergarten and second grade (*p* = 0.06) (see Figure 5, right). We did not find any group main effect in condition 1 (W–PF, coarse print tuning effect, reading group: all *F*(2,47) < 1.02, *p* > 0.36; grade group: all *F*(2,47) < 1.08, *p* > 0.35)) or condition 3 (OLPW–OINW, sublexical fine tuning effect, reading group: *F*(2, 47) = 0.58, *p* = 0.56; grade group: *F*(2, 47) = 2.05, *p* = 0.14). However, in condition 3, we found a variation in the spatial filter topographies. Inspection of the topographies shown in Figure 6A reveals a lateralization shift in condition 3 (OLPW–OIPW), with right-lateralized visual occipital activation in kindergarten children shifting to left lateralization in first and second graders. In contrast, as shown in Figure 6B, no topographical difference was revealed across grade levels in condition 1 (W–PF). Topographies of condition 2 (W–OLPW) are not presented here as the first RCs did not correspond to statistically significant responses for all grade groups.

**Figure 5:**
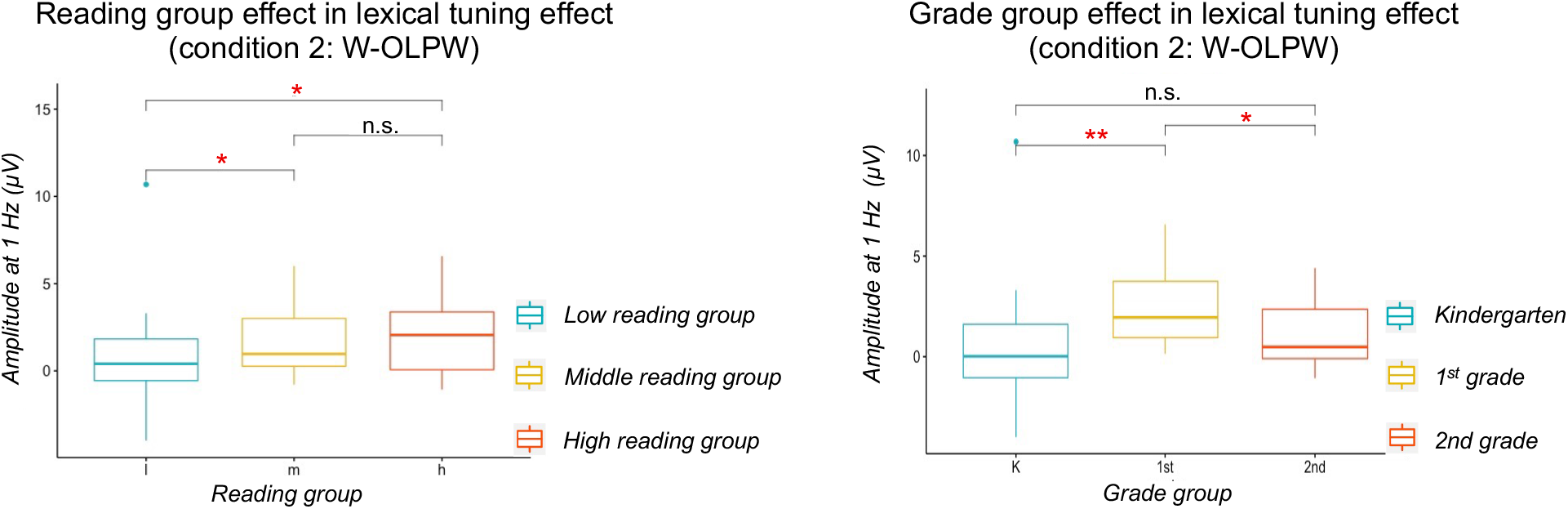
Group main effects in lexical fine tuning (condition 2: W–OLPW) at beginning readers. Reading group (left) and grade group (right) comparisons of projected amplitude at the first significant harmonic of deviant frequency (1 Hz). Left panel: response amplitudes were stronger in middle and high reading groups than in the low reading group, no significant difference was found between middle and high groups. Right panel: response amplitudes were stronger in first grade than in kindergarten and second grade, marginal significant difference was found between kindergarten and second grade, **: *p* < 0.01, *: *p* < 0.05, n.s.: not significant.

**Figure 6:**
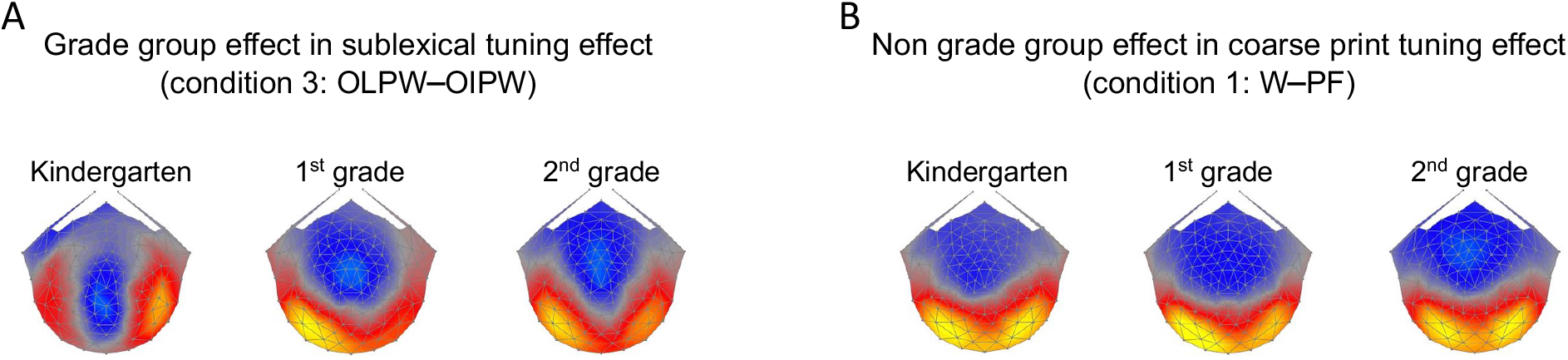
Grade main effects in sublexical fine tuning (condition 3: W–OLPW) at beginning readers. A: right to left lateralization shift in sublexical fine tuning effect: left lateralized topography map in Kindergarten (left), right lateralized topography maps in first grade (middle) and second grade (right). B: Consistent bilateral topography maps in coarse print tuning effect (condition 1: W–PF): topography maps in Kindergarten (left), right lateralized topography maps in first grade (middle) and second grade (right).

### 3.2 Base Analyses Results

For the base frequency and its harmonics, RCA was computed on sensor-space data across three conditions together in order to facilitate direct quantitative comparison of the projected data in a shared component space. The results are summarized in Figure 7. Panel A shows topographic maps of the spatial filters for RCs 1, 2, and 3, respectively. The RC1 resulting from all conditions trained together is distributed over occipito-temporal regions, RC2 is focally distributed over the primary visual cortex, and RC3 is centered on medial occipital area. The summary plots of the responses in the complex plane (Panel B), show overlapping amplitudes (vector lengths) and phases (vector angles) across three conditions. As shown in Panel C, response amplitudes did not differ significantly across the five even harmonics between 2 Hz and 10 Hz (ANOVA, RC1: *F*(2,143) ≤ 0.94, *p* ≥ 0.39; RC2: *F*(2,143) ≤ 1.66, *p* ≥ 0.19; RC3: *F*(2,143) ≤ 0.39, *p* ≥ 0.67). Derived latency estimations for each component were also similar across conditions (RC1: 167.37 ms, 174.80 ms, and 168.00 ms for conditions 1, 2, and 3 respectively; RC2: 285.38 ms, 281.22 ms, and 280.34 ms for conditions 1, 2, and 3 respectively; RC3: 330.97 ms, 329.13 ms, and 328.85 ms for conditions 1, 2, and 3 respectively). Thus, we consider the responses at the base frequency to be highly comparable across the three conditions.

**Figure 7:**
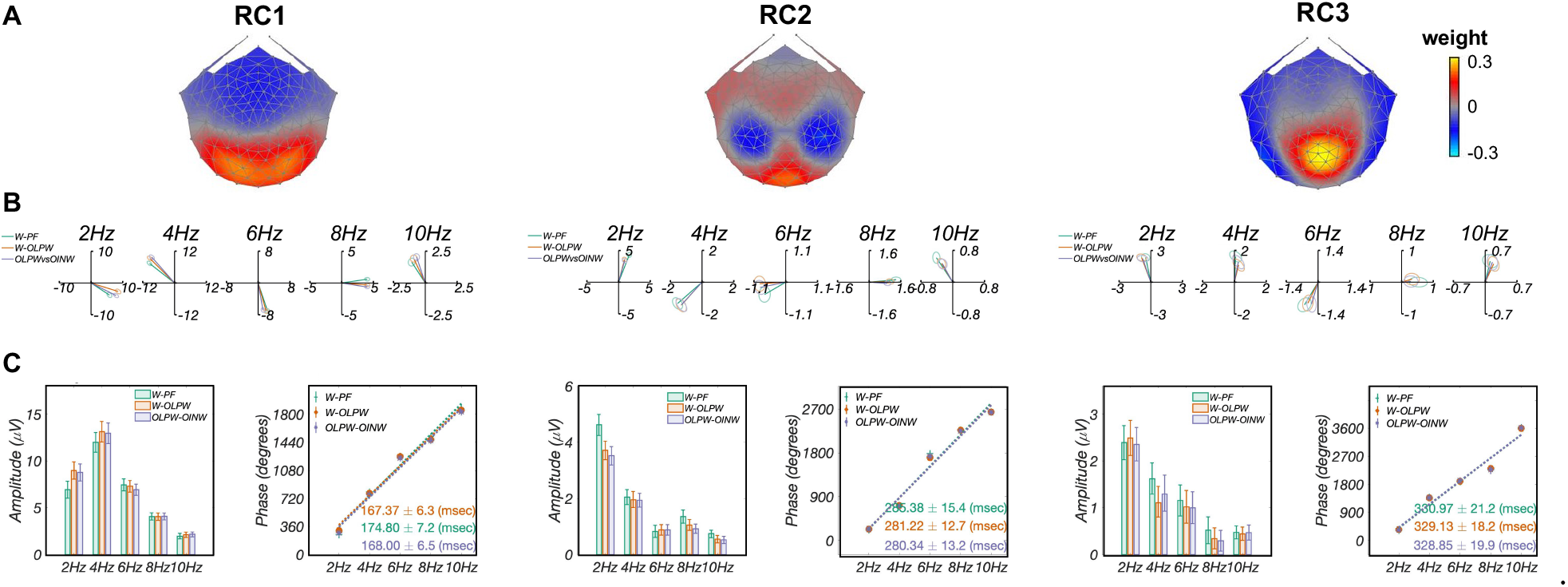
Base analysis comparisons across three conditions. A: Topographic visualizations of the spatial filters for the first three components: RC1, RC2, and RC3; B: Response data presented in the complex plane, where amplitude information is represented as the length of the vectors, and phase information in the angle of the vector relative to 0 degree (counterclockwise from 3 o’clock direction), ellipse indicates standard error of the mean (*SEM*); C: Comparison of projected amplitude (left) and latency estimation (right) across three conditions. Response amplitudes did not differ significantly across the five harmonics (RC1: *p* > 0.75; RC2: *p* > 0.19; RC3: *p* > 0.67), latency estimation derived from phase slopes across harmonics were also similar across conditions (RC1: 167.37 ms, 174.80 ms, and 168.00 ms for conditions 1, 2, and 3 respectively; RC2: 285.38 ms, 281.22, and 280.34 ms for conditions 1, 2, and 3 respectively; RC3: 330.97 ms, 329.13, and 328.85 ms for conditions 1, 2, and 3 respectively).

### 3.3 Behavioral Results

For the 3-in-a-row repetition detection task, *d*’ were *m* = 1.10, *s* = 0.18, *m* = 1.17, *s* = 0.22, *m* = 1.08, *s* = 0.29 for conditions 1, 2, and 3, respectively. A one-way ANOVA showed that *d*’ did not differ significantly across the three conditions (*F*(2,141) = 0.44, *p* = 0.64). Thus, we conclude that participants were equally engaged throughout the three conditions of the experiment.

## 4 Discussion

In this study, we separated the cognitive processes and neural mechanisms underlying coarse print tuning from sublexical orthographic regularity and lexicality effects involved in word recognition in beginning readers. Combining a Steady-State Visual Evoked Potentials (SSVEPs) approach with Reliable Components Analysis (RCA), we found, for the first time, robust responses to both sublexical and lexical representations, even in kindergarteners. In addition, sublexical fine tuning depended on age, with right-lateralized responses in kindergarteners that shifted leftward in later grades. Lexical tuning was related to reading fluency, with lower-amplitude responses in the low reading fluency group than in middle and high reading groups. Finally, in line with recent studies with adults (Wang et al., 2021; Woolnough et al., 2021; Lerma-Usabiaga et al., 2018), beginning readers also have multiple neural sources involved in coarse print tuning.

### 4.1 Comparison between component-space analyses and sensor-space analyses

Conventional procedures for analyzing ERP (e.g., Maurer et al. (2006)) and SSVEP (e.g., Lochy et al. (2015)) data involve preselection of one or a few literature- or SNR-based sensors in a cluster and then analysis of the evoked responses in that chosen spatial subset. There are two limitations of this approach. First, it has been suggested that sensor selection should not be based on where the effect of interest is strongest, because this procedure leads to a bias in reporting false positives (Kilner, 2013). Second, a single region of interest (ROI) cannot separately describe activity in multiple underlying sources. Multiple brain regions are likely to be functionally and structurally connected with each other during higher-level neural processes. For instance, middle and anterior vOT is connected with other language areas (e.g., the angular gyrus, supramarginal gyri and the superior temporal gyrus), allowing orthographic-lexical-semantic transformations to occur (Lerma-Usabiaga et al., 2018; Woolnough et al., 2021). A single subset of sensors is unlikely to capture and isolate different neural processes arising from different underlying cortical sources that have separate topographies, as these would need to be characterized by multiple ROIs.

In contrast, spatial filtering approaches compute weighted linear combinations across the whole montage of sensors, yielding signal “components” that can capture the presence of multiple underlying sources in the data. Among other spatial filtering approaches, such as Principal Components Analysis (PCA) and Common Spatial Patterns (CSP) (Blankertz et al., 2008), RCA achieves higher SNR with lower trial count by maximizing across-trial correlations (i.e., “reliability”) while minimizing noise power (Dmochowski et al., 2015). It has been shown that the topographies of components derived from RCA more strongly resemble the underlying lead fields generating the observed SSVEPs than PCA and CSP (Dmochowski et al., 2015). Thus, the topographies of components inform the (approximate) locations of the underlying cortical sources. In addition, given the assumption of constant response phases over repeated trials in the SSVEP context, the component optimization procedure reflects the underlying phase-locked activity of the SSVEP (Norcia et al., 2015). Moreover, by measuring the change of phase over the different response frequencies with significant signals, the slope of this relationship can be interpreted as a response latency (Norcia et al., 2020). Latency estimates of one or more components can provide insight into temporal dynamics and processing order of different cortical sources, extending the majority of previous SSVEP studies that have focused on topographies and response amplitudes.

### 4.2 Three distinct sources and processing times underlie coarse print tuning in beginning readers

RCA of word deviants in the word–pseudofont (W–PF) contrast produced three components with differing topographies and time courses (see Figure 3).

More importantly, we find that coarse print tuning (i.e., W vs. PF) is already observed in kindergarteners who have received only 2.5 months of school reading education on average. This is consistent with a recent SSVEP study which found specialization for letter strings in preschoolers, suggesting that coarse print tuning emerges earlier than usually thought (Lochy et al., 2016). In previous ERP studies, coarse print tuning has typically been detected only after 1–1.5 years of formal reading education (Eberhard-Moscicka et al., 2015; Maurer, Brem, et al., 2005; Maurer et al., 2006). The early emergence of coarse print tuning in the current study and in the previous study by Lochy et al. (2016) may be partially due to the SSVEP paradigm used. The SSVEP paradigm differs from transient ERP approaches having lower signal to noise ratio that might thus be unable to detect this print specialization, especially in young readers. Overall, the early emergence of coarse print tuning suggests that only limited knowledge of perceptual information associated with letters and words is sufficient to develop underlying circuits that are specialized for print processing (Chyl et al., 2018). This view is supported by training studies that have shown print sensitivity to emerge rapidly even after a short duration (i.e., hours) of reading training in young, non-reading kindergarten children (Brem et al., 2009; Pleisch et al., 2019).

#### 4.2.1 RC1: left vOT activation at 300 ms

Our finding of preferential responses to W vs. PF in the left vOT (RC1) is in agreement with many previous fMRI studies (e.g., adolescents (Brem et al., 2005); 10.3-year-old children (Brem et al., 2009); 7–14-year-old children (Centanni et al., 2017)) and with recent EEG studies using SSVEP paradigms (adults (Lochy et al., 2015); preschoolers and 1st and 2nd graders (Lochy et al., 2016; van de Walle de Ghelcke et al., 2021)). The left vOT—also referred to as the visual word form system (VWFS)—plays a crucial role in fluent and efficient reading (Baker et al., 2007; McCandliss et al., 2003; Dehaene & Cohen, 2011). Within the VWFS, a word form sensitive area, the “visual word form area” (VWFA), which is centered in the left mid fusiform gyrus, is particularly sensitive to print (Centanni et al., 2017; Dehaene et al., 2005; Vinckier et al., 2007; Glezer & Riesenhuber, 2013) and whole visual word forms (Binder et al., 2006; Glezer et al., 2009).

The coarse sensitivity of vOT/VWFA to print has been linked to domain-specific visual perceptual expertise that arises from experience with words and/or letter strings during reading acquisition, rather than merely general maturation which would affect both words and pseudofonts equally (Brem et al., 2005; Maurer et al., 2006; McCandliss & Noble, 2003; B. A. Shaywitz et al., 2002). Visual experience and the acquisition of grapheme-phoneme correspondences for print drive a functional reorganization in the response properties of the VWFA to become specifically tuned to recurring properties and structural regularities of a writing system.

Phase-lag quantification of RC1 provides a latency estimate of around 300 ms. This 300 ms latency is consistent with previous pediatric ERP studies showing that print sensitivity occurs at around 180–300 ms (Maurer, Brem, et al., 2005), slightly later than that in adults at around 140–200 ms (Brem et al., 2005; Maurer et al., 2006; Eberhard-Moscicka et al., 2015; Wang & Maurer, 2020), which possibly reflects additional processing energy due to lack of visual familiarity with words or letter knowledge in younger children (Maurer, Brandeis, & McCandliss, 2005).

#### 4.2.2 RC2: left central-parietal activation at 350 ms

In addition to the left vOT activation (RC1), we also observed a component that was maximal over left central-parietal regions (RC2). A linear fit across phases at significant harmonics revealed a latency estimate of around 350 ms. This 350 ms latency is consistent with EEG (Grainger et al., 2006; Carreiras et al., 2009) and MEG (Pylkkänen & Marantz, 2003; Fruchter & Marantz, 2015) results of adult studies showing that lexical access arises later than initial form-based processing. For example, Carreiras et al. (2009) found phonology effects at 350–550 ms post-stimulus onset.

We consider preferential word responses measured over left central-parietal electrodes (RC2) as most likely implicating access of lexical-semantic representations. This interpretation is consistent with accumulating evidence showing that other higher-level linguistic representation regions are also involved in visual word recognition through functional and/or structural connections with left vOT (Lerma-Usabiaga et al., 2018; Yeatman et al., 2013; Seghier & Price, 2013; White et al., 2019; Kay & Yeatman, 2016). For example, recent fMRI studies in both adults and children have shown strong functional connectivity between VWFA and superior temporal and inferior parietal regions (Li et al., 2017; Stevens et al., 2017; Chen et al., 2019), which have been implicated in phonological processing (Jobard et al., 2003). Therefore, the left central-parietal brain activation in our study may reflect grapheme-to-phoneme conversion and/or accessing of lexical semantic information (Woolnough et al., 2021), by iterating feed-forward and feedback signal from and to left vOT (Price & Devlin, 2011).

#### 4.2.3 RC3: activation over early visual cortex at 320 ms

Finally, a third component that is maximal focally over early visual cortex (RC3) was found. This result is in line with several previous studies with adult participants, and the notion that reading expertise also relies on perceptual learning in earlier levels of the visual system than the VWFA (Szwed et al., 2011, 2014; Nazir, 2000; Nazir et al., 2004; Nazir & Huckauf, 2007). For example, Nazir and colleagues found more activation for intact words than for scrambled words in early visual areas V1/V2 and V3/V4 (Nazir, 2000; Nazir et al., 2004). In addition, this visual specialization in primary visual cortex is more prominent for reading than for objects, as no difference was found in this area between intact and scrambled objects (Nazir, 2000; Nazir et al., 2004; Nazir & Huckauf, 2007). Increased sensitivity of early visual and occipital cortex to words might be driven by the perceptual learning mechanisms underlying reading (Szwed et al., 2011, 2014). In contrast to visual processing of objects with varied locations and sizes, the unique style of rapid, high spatial frequency, and massively parallel processing of letters and words may place a strong constraint on early visual processing, which leads to plastic changes in early visual cortex (Dehaene et al., 2015; Li et al., 2017; Poort et al., 2015).

The 320 ms latency of RC3 may reflect feedback projections to early visual cortex from higher-level visual (e.g., VWFA) and language areas involved in linguistic and phonological processing (Wheat et al., 2010; Barzegaran & Norcia, 2020), in combination with bottom-up mechanisms from this area back to VWFA (Church et al., 2011).

To summarize, three components were derived from responses to condition 1 (W–PF) through a data-driven analysis of SSVEP. The current findings provide evidence for the interactive account model in which visual word recognition involves recurrent interactions between early visual cortex, VWFA, and lexical/phonological areas (Price & Devlin, 2011). However, these interpretations are speculative without the support of functional connectivity and source localization data. Thus, the exact recurrent mechanisms of different levels of the visual system and language areas need to be verified in future studies.

### 4.3 Lexicality effects emerge in kindergarten and correlate with reading efficiency and grade in beginning readers

RCA of word deviants in the words-pseudowords contrast generates a component that is broadly distributed over occipital-temporal/parietal regions. It is important to note that the current study has for the first time found a lexical fine tuning effect even in kindergarten children who averaged 5.9 years of age, a younger population than has been reported in previous transient ERP studies. For example, Maurer and colleagues found stronger responses for words than for pseudowords in second grade children (Maurer et al., 2006), but not in non-reading kindergarten children (Maurer, Brandeis, & McCandliss, 2005), and not in adults (Maurer, Brem, et al., 2005). By contrast, Posner & McCandliss (1999) found that fine tuning for words was not fully developed even in 10-year-olds.

Our detection of a lexicality effect in kindergarteners may be due to several distinctive aspects of the present study. First, very high frequency (on average 2110/million) 3-letter words were used, instead of 5-letter words with relatively low frequency (on average 79/million) in another study (Zhao et al., 2014). By recruiting a large group of 7-year-olds (first grade) with wide range in reading ability, Zhao and colleagues found lexical fine tuning for words only in children with high reading ability, but not in those with low reading ability. Second, the processing demands of the 3-in-a-row repetition detection task in the current study are higher than the implicit tasks used in other studies (for instance, color detection task in Zhao et al. (2014); fixation cross color change task in Lochy et al. (2016)). As a result, higher-level reading-related processes (herein lexical processing) may have been better engaged in the present study. Finally, it has been suggested that the effect size of the lexicality effect is rather small and therefore it may not always be reliably detected (Eberhard-Moscicka et al., 2015). The SSVEP paradigm we used is more sensitive toward detecting discrimination responses because the noise at the paradigm’s predefined and specific frequencies is smaller than the noise in transient ERP paradigms—which is broadband, as is the response itself (Lochy et al., 2016). Despite this, using a similar SSVEP paradigm, Lochy et al. (2016, 2020) did not find a W-PW difference in either kindergarteners or in first and second graders. This may be because of the fast presentation rates (1.2 Hz and 6 Hz) used in Lochy and colleagues’ studies, while we used 1 Hz and 2 Hz, given evidence that a slower alternation presentation mode elicits responses with higher signal-to-noise ratio (Wang et al., 2021; Yeatman & Norcia, 2016).

The observed activation over occipital-temporal/parietal electrodes in the present study is in line with previous fMRI studies that have shown that anterior occipital-temporal regions are engaged in lexical or semantic processing, whereas posterior occipito-temporal regions are more associated with pre-lexical form-based processing (Vinckier et al., 2007; Lerma-Usabiaga et al., 2018). As an example, a recent intra-cranial study found two significant clusters—the mid-fusiform cortex and lateral occipitotemporal gyrus—in a word–pseudoword contrast, with lateral occipitotemporal cortex displaying sensitivity to sublexical and lexical processing later than mid-fusiform cortex (Woolnough et al., 2021). Similarly, a combined EEG/MEG study also found more activation at 180 ms to words compared to pseudowords in the left anterior middle temporal lobe, which was interpreted as early lexico-semantic retrieval (Hauk et al., 2012).

In addition to finding an early emergence of lexical fine tuning in kindergartners, we further found that this lexical tuning is influenced by word reading efficiency and grade. Larger amplitudes for word deviants were found in middle and high reading fluency groups than in the low reading group, as well as in first grade than in kindergarten and second grade. Reading group effect may reflect different levels of automaticity in higher-order linguistic processes in children with different reading outcomes (Roembke et al., 2019). For children who read words frequently, the orthographic-lexical-semantic transformation is more automatized, and thus, higher levels of reading-related processes maybe involved during brief stimulus presentations (Zhao et al., 2014). This may be reflected by larger amplitudes as in the current study or stronger brain activation as in fMRI (e.g., Hauk et al. (2012)). In relation to grade group differences—stronger amplitude in first grade than in kindergarten and second grade—of lexical tuning, an inverted U-shaped developmental trajectory of coarse print tuning was found in a previous ERP study, with specialization for print peaking after learning to read in less than two years and then decreasing with further reading experience in adults (Maurer et al., 2006). However, neither a group-level effect of grade nor a correlation with reading efficiency were found in the discrimination responses between words and pseudofonts (coarse tuning). This is consistent with the notion that fine tuning, instead of coarse tuning, might be a bio-marker of reading ability at the initial stage of learning to read (Zhao et al., 2014).

To summarize, the present finding of a lexicality effect as early as kindergarten contrasts with previous ERP studies, which suggests that fine tuning for words develops slowly over the course of reading acquisition. With more sensitive measures, the fine tuning for words—as a bio-marker of reading ability—can be detected at a much earlier stage than previously assumed.

### 4.4 Right to left occipital topographic shift of sublexical orthographic regularity in beginning readers

The spatial component reflecting the orthographic regularity effect, indicated by discriminative responses to orthographically legal pseudowords (OLPW) versus orthographically illegal nonwords (OINW) manifests as a scalp topography that is distributed over left occipital regions. Left occipital regions have been previously shown to be sensitive to processing of orthographic representations (Samuelsson, 2000). Our cross-sectional analysis over grade level showed a right-to-left lateralization shift from kindergarten to first and second grades. Orthographic regularity effects in left occipital regions have also been found in previous fMRI (Woollams et al., 2011; Samuelsson, 2000; Turkeltaub et al., 2003) and transient ERP studies involving both children (Zhao et al., 2014) and adults (Proverbio et al., 2004; Araújo et al., 2015; Hauk et al., 2006). For instance, Araújo et al. (2015) found significant larger negativities to pseudowords than to nonwords at left occipital electrodes, but only in normal reading adults and not in a dyslexia group. Developmentally, Zhao et al. (2014) found that EEG response differences between orthographically legal pseudowords and consonant strings over the left hemisphere (occipito-temporal area) correlated with reading speed in 7-year-old children, with faster reading speed in children showing stronger responses to pseudowords relative to consonant strings.

Our finding that the sublexical fine tuning effect (condition 3: OLPW–OINW) can be demonstrated as early in kindergarten supports the notion that readers can very quickly decode new orthographic regularities, even in fully unfamiliar scripts (Chetail, 2017). In addition, other studies have suggested that sublexical orthographic processing occurs at a very early stage (80–120 ms) of the reading process (Coch & Mitra, 2010; Hauk et al., 2006; Araújo et al., 2015). As a prerequisite for reading, sensitivity of orthographic regularities can facilitate access to phonological and morphemic structure of words (Chetail et al., 2015). Thus, measuring sublexical and lexical orthographic processing, as in the current study, is helpful for better understanding how children’s brains develop different levels of visual-orthographic processing.

Our results add to previous studies by showing the right-to-left hemisphere shift from kindergarten to first and second grades. This lateralization shift most likely represents changes in processing strategy during reading acquisition. This interpretation is in line with the finding that unlike the left hemisphere, which remains important over the course of reading acquisition, right hemisphere non-lexical form recognition systems are progressively disengaged (Maurer & McCandliss, 2007; S. E. Shaywitz & Shaywitz, 2007; Turkeltaub et al., 2003; Pleisch et al., 2019). Therefore, kindergarten children with limited orthographic knowledge may rely more on a non-lexical recognition system in the right hemisphere, and then progressively shift to using the left hemisphere after accumulating knowledge of letters and words. Precise measurements of the neural and behavioral changes within individuals through a longitudinal study are needed in the future to establish a causal role for the shift.

### 4.5 Low-level visual feature processing during word recognition

Unlike previous SSVEP work on base rate responses (Lochy et al., 2015, 2016, 2020) which has only focused on medial occipital sites (around electrode Oz), three distinct components were derived from base stimulation responses using RCA. The first component is distributed over ventral occipito-temporal cortex. Occipito-temporal regions have been linked to processing of learned visual feature information associated with letter (Barzegaran & Norcia, 2020; Wong et al., 2005) and word forms (McCandliss et al., 1997, 2003). The second component is maximal focally at primary visual cortex. Early/primary visual cortex receives visual input, including of objects and words, from the eyes through the lateral geniculate nucleus (LGN, Sadato et al. (1996)). Finally, a component centered on medial occipital electrodes was also revealed. Medial occipital electrodes have been shown to support early stages of visual processing, i.e., low-level visual features analysis, including size, shape, configuration, line junctions and letter segments (Ostwald et al., 2008; Dehaene et al., 2005).

Using the same RCA approach, our recent study with adult participants found only one significant component (similar to the RC3 found here) underlying base stimulation (Figure 2 in Wang et al. (2021)). In addition to the age difference of participants, a main difference between these two studies is their stimulus presentation rates, with 6-Hz vs. 2 Hz base frequencies in the adult study and the current study, respectively. Thus, it is possible that more information from the base stream can be processed due to the longer duration of each stimulus at the slower presentation rate.

Overall, the three base-response related components of the current study may reflect stages in the pathway for visual recognition that progress from basic visual input perception in primary visual cortex to processing of line junctions and letter fragments in medial occipital area, and then to processing in ventral occipio-temporal areas for pre-lexical letter and word form recognition (Dehaene et al., 2005; Dehaene & Cohen, 2011). Basic low-level visual features (e.g., luminance, size, shape, letter fragments) were well matched across four different types of stimuli (W, PF, OLPW, OINW). This feature-matching could explain comparable responses, including phase, amplitude, and topographies, across different contrasts.

## 5 Conclusion

In conclusion, the current study examined letter-selectivity, orthographic regularity, and lexicality effects in beginning readers using a SSVEP paradigm and a data-driven component approach (RCA). Three functionally distinct processes with different temporal dynamics were found to underpin coarse print tuning (letter-selectivity). Sensitivity to lexical content emerged as early as in kindergarteners and was associated with reading efficiency. Furthermore, sensitivity to English language orthography was also present starting from kindergarten in source regions that later shifted from right-to left-hemisphere electrodes. Our results provide new insights on the functional circuits underlying print tuning and revealed different and robust brain activations for sublexical and lexical orthographic processing in very early readers.

## Abbreviations

(RCA): Reliable Components Analysis
(RC1): Reliable Component 1
(RC2): Reliable Component 2
(RC3): Reliable Component 3
(SSVEP): steady-state visual evoked potentials

## Acknowledgments

We thank the students, their families, and teachers for participating. We also thank Rachana Pillai for her help with data collection.

